# Genetic incompatibilities in reciprocal hybrids between populations of *Tigriopus californicus* with low to moderate mitochondrial sequence divergence

**DOI:** 10.1101/2022.09.19.508600

**Authors:** Timothy M. Healy, Ronald S. Burton

## Abstract

All mitochondrial-encoded proteins and RNAs function through interactions with nuclear-encoded proteins. These interactions are critical for mitochondrial function and eukaryotic fitness, and coevolution maintains inter-genomic (i.e., mitonuclear) compatibility within taxa. Hybridization can disrupt coevolved interactions, resulting in hybrid breakdown, and mitonuclear incompatibilities may be important mechanisms underlying reproductive isolation and, potentially, speciation. Recently, signatures of strong selection to maintain compatible mitonuclear genotypes in hybrids have been detected in at least some inter-population crosses. However, this work has only been conducted in crosses between populations with extremely high levels of genetic divergence in both their mitochondrial and nuclear genomes, leaving the generality of strong selection for mitonuclear compatibility unclear. Here we address this limitation with reciprocal inter-population F_2_ hybrids between relatively low-divergence populations of the intertidal copepod *Tigriopus californicus*. Our results show that the dominance of mitonuclear effects consistent with coevolved mitonuclear genotypes in fast-developing (i.e., high-fitness) hybrids is reduced in low-divergence crosses, but that selection to maintain mitonuclear compatibility is still observed on some nuclear chromosomes. Consequently, we demonstrate that, even at low levels of genetic divergence between taxa, mitonuclear incompatibilities may play a key role in the early stages of reproductive isolation.

## Introduction

Mitochondrial dysfunction can lead to dramatic reductions in organismal performance and fitness as oxidative phosphorylation supplies the majority of cellular energy in eukaryotes through the generation of ATP (e.g., Lane 2005). Mitochondrial functions are highly reliant on interactions between proteins and RNAs encoded in the mitochondrial and nuclear genomes, and compatibility between gene products of the two genomes is typically maintained by coevolution within independent taxa (Rand et al. 2004; Burton et al. 2013; Hill 2015). However, hybridization between populations or species breaks apart genes subject to inter-genomic coevolution, because in second- and higher-generation hybrids, mitochondrial genotypes are found on nuclear backgrounds with regions that are homozygous for foreign nuclear alleles (Burton et al. 2006). Thus, given the overall links between mitochondrial performance and fitness, mitonuclear incompatibilities are potentially key mechanisms underlying hybrid breakdown, reproductive isolation and speciation (Gershoni et al. 2009; Burton and Barreto 2012; Hill 2016; Hill et al. 2019), particularly at early stages of isolation due to the relatively high evolutionary rate of the mitochondrial genome (Lynch 1997; Wallace 2010; Burton and Barreto 2012).

For mitonuclear compatibility to play a substantial role in reproductive isolation, there would have to be strong selection favouring compatible genotypes among hybrid organisms (Sloan et al. 2017; Hill et al. 2019), such that naturally occurring hybrids carrying incompatible mitonuclear genotypes would be unlikely to persist. Yet, despite known fitness consequences of mitonuclear incompatibilities in several taxa (e.g., Ellison and Burton, 2008b; Meiklejohn et al. 2013) and some evidence for strong selection associated with variation among hybrids (Healy and Burton 2020; Han and Barreto 2021), introgression of mitochondrial genotypes across species or population boundaries has been observed in many cases (Chan and Levin 2005; Toews and Brelsford 2012; Sloan et al. 2017), and mitonuclear incompatibilities in humans (*Homo sapiens*) are not typically major concerns in medical contexts such as mitochondrial replacement therapies (Eyre-Walker 2017). As a result, the extent to which selection for inter-genomic compatibility presents a substantial barrier for gene flow among taxa in general remains largely unresolved (Hill 2016, 2019; Sloan et al. 2017; Burton 2022).

Evidence for strong selection associated with mitonuclear interactions comes from studies on inter-population hybrids of the marine copepod *Tigriopus californicus* (Healy and Burton 2020; Han and Barreto 2021). This species inhabits intertidal splash pools on the Pacific coast of North America from Baja California, Mexico to Alaska, USA with substantial genetic divergence among populations from different rocky outcrops (e.g., Burton and Lee 1994; Edmands 2001; Burton et al. 2007; Barreto et al. 2018). Although inter-population hybrids are viable in the laboratory, loss of fitness as a result of mitonuclear incompatibilities has been consistently observed across many *T. californicus* studies (Edmands and Burton 1999; Harrison and Burton 2006; Ellison and Burton 2006, 2008b; Healy and Burton 2020; Han and Barreto 2021; Pereira et al. 2021). In particular, variation in developmental rate among F_2_ hybrids has been associated with mitochondrial performance and the degree of mitonuclear compatibility in crosses between a population from San Diego, California and populations from Santa Cruz California (Healy and Burton 2020) or Strawberry Hill Wayside, Oregon (Han and Barreto 2021). In both of these cases, there are high levels of genetic divergence between the parental populations contributing to these crosses. For instance, the sequences of the San Diego and Santa Cruz mitochondrial genomes are 21% divergent overall (Barreto et al. 2018), and the sequences of the mitochondrial-encoded protein-coding genes differ by 11% to 35% between the San Diego and Strawberry Hill Wayside populations (Han and Barreto 2021). Similarly, median sequence divergences across nuclear-encoded genes between copepods from San Diego and Santa Cruz or Strawberry Hill Wayside are 2.5% or 2.8%, respectively (Han and Barreto 2021). Consequently, the extent to which the evidence for strong effects of mitonuclear interactions in *T. californicus* can be generalized to other species is unclear. For instance, in *Drosophila sp.*, which are another well-established model for the study for mitonuclear interactions (Dowling et al. 2007; Hoekstra et al. 2013; Meiklejohn et al. 2013; Carnegie et al. 2021; Rand et al. 2022), variation in mitochondrial genome sequences even among species is much lower than among *T. californicus* populations (e.g., ~4.5% between *D. melanogaster* and *D. simulans*; Ballard 2000). *T. californicus* populations offer an ideal opportunity to address this issue, because the amount of genetic divergence among populations is variable across the species range (Edmands 2001; Peterson et al. 2013; Pereira et al. 2016) and the phenotypic effects of mitonuclear interactions demonstrate wide ranges of impact among crosses between different populations (Burton 1990).

In the current study, we examine variation in nuclear allele frequencies associated with differences in developmental rate among reciprocal F_2_ hybrids between two pairs of *T. californicus* populations: San Diego and Bird Rock in southern California, USA, and Santa Cruz and Pescadero Beach in central California. These population pairs have relatively low levels of inter-population genetic divergence (Pereira et al. 2016), allowing selection among hybrids associated with mitonuclear compatibility to be assessed at less extreme degrees of sequence divergence than in past studies (Healy and Burton 2020; Han and Barreto 2021). In particular, here we assess allele frequency patterns consistent with mitonuclear and nuclear-only effects on variation in developmental rate (a proxy for fitness) within and between the reciprocal crosses, and we use these data to evaluate the potential consequences of mitonuclear incompatibilities at low levels of genetic divergence.

## Materials and methods

### Population sampling and copepod husbandry

Adult *T. californicus* were collected from four locations along the coast of California, USA in the spring of 2018: San Diego (SD: 32° 44′ 45″ N, 117° 15′ 18″ W), Bird Rock (BR: 32° 48′ 51″ N, 117° 16′ 24″ W), Santa Cruz (SC: 36° 56′ 58″ N, 122° 02′ 49″ W) and Pescadero Beach (PE: 37° 15′ 35″ N, 122° 24′ 51″ W). At each site, copepods were removed from supralittoral tidepools with large plastic pipettes, and transferred into 1 L plastic bottles containing water from the tidepools. The copepods were transported to Scripps Institution of Oceanography, University of California San Diego within 24 h of collection, and were established in population-specific laboratory cultures of 250 mL filtered seawater (0.44 μm pore size; 35 psu) held in 400 mL glass beakers that were placed inside incubators set to 20 °C and 12h:12h light:dark. Copepods consumed natural algal growth within the cultures, and were also fed once weekly with powdered spirulina (Salt Creek, Inc., South Salt Lake City, UT, USA) and ground TetraMin® Tropical Flakes (Spectrum Brands Pet LLC, Blacksburg, VA, USA). These cultures and conditions were maintained for at least one month (~1 generation) prior to initiating experiments.

### Experimental crosses and classification of fast- or slow-developing F_2_ hybrids

Virgin *T. californicus* females from each population were obtained by splitting mate-guarding pairs (Burton 1985), and were used in two sets of reciprocal inter-population hybrid crosses (4 crosses total): SD♀ x BR♂ (SDxBR), BR♀ x SD♂ (BRxSD), SC♀ x PE♂ (SCxPE) and PE♀ x SC♂ (PExSC). Note that the reciprocal crosses have equivalent hybrid nuclear genomes under neutral expectations, whereas the mitochondrial genome is inherited from the maternal population (e.g., Lima et al. 2019). Crosses were performed by placing 40 virgin females of a desired population in a 2 × 10 cm petri dish containing ~60 mL of filtered seawater, and then adding 40 males from the other population in the cross. This was repeated three times per cross, such that 120 total matings were performed in each case. The females and males paired haphazardly within the dishes, and gravid females were transferred to a new dish once they were observed. F_1_ offspring hatched naturally, and once the F_1_ hybrids were visible to the naked eye, the P0 females were removed. F_1_ females and males mated haphazardly, and gravid F_1_ females were transferred to new dishes when they were observed. Mature (i.e., red) egg sacs were chosen haphazardly for the F_2_ developmental trials.

Variation in developmental rate among F_2_ hybrids was assessed as described in Healy and Burton (2020). In brief, two weeks after the initial transfers of gravid F_1_ females to their new dishes, mature F_2_ egg sacs were dissected from the females and placed into wells of Falcon® 6-well plates containing filtered seawater (≤ 4 egg sacs per well). F_2_ offspring hatched overnight, and were fed with powdered spirulina as described above. During development in *Tigriopus sp.*, there is a distinct metamorphosis from the last naupliar stage (N6) to the first copepodid stage (C1; e.g., Raisuddin et al. 2007), and the time to this metamorphosis can be used as a measure of developmental rate. Healy and Burton (2020) scored developmental rate for individual SDxSC and SCxSD F_2_ hybrids, and defined fast- or slow-developing copepodids as those that metamorphosed 8-12 or ≥ 22 days post hatch (dph), respectively. In the current study, we used the same dph to metamorphosis to define fast and slow developers in our SDxBR, BRxSD, SCxPE and PExSC crosses; however, developmental rates were not scored for individual copepodids. Instead, fast-developing groups were established by transferring all copepodids present in the 6-well plates at 12 dph to new cross-specific petri dishes, and slow-developing groups were established by discarding all copepodids present in the 6-well plates at 21 dph and then transferring all remaining nauplii to new cross-specific dishes. Between 83 and 221 F_2_ egg sacs per cross were assessed in the developmental rates trials.

### Pool-seq: DNA isolation and sequencing

Once the fast- and slow-developing F_2_ hybrids reached adulthood, approximately 160 haphazardly selected copepods from each group were pooled and preserved at −80 °C. DNA was isolated from the frozen pools of individuals by phenol-chloroform extraction (Sambrook and Russell 2006; as described in Healy and Burton 2020). In brief, the pools were homogenized by hand in 150 μL of buffer (200 mmol L^−1^ sucrose, 100 mmol L^−1^ NaCl, 100 mmol L^−1^ Tris-HCl pH 9.1, 50 mmol L^−1^ EDTA, 0.5% SDS) and digested at 56 °C overnight with 100 μg of Proteinase K (Thermo Fisher Scientific, Waltham, MA, USA) in a total buffer volume of 400 μL. The digests were treated with 25 μg RNaseA (Thermo Fisher Scientific), and then 200 μL of 5 mol L^−1^ potassium acetate was added. After 10 min on ice, the digests were centrifuged at 13,000 g for 10 min, supernatants were obtained, 400 μL each of buffer-saturated phenol (Thermo Fisher Scientific) and chloroform (EMD Millipore Corporation, Darmstadt, Germany) were added to the supernatants, and the mixtures were centrifuged at 20,000 g for 5 min. Aqueous phases were transferred to new tubes, and organic phases were extracted again with 400 μL chloroform followed by centrifugation at 15,000 g for 1 min. The second set of aqueous phases were again transferred to new tubes, and all aqueous phases had 1,200 μL of ice-cold 95% ethanol added prior to incubation at −20 °C for 1 h. Precipitated DNA was collected by centrifugation at 16,000 g for 20 min. The two DNA pellets for each sample were pooled and washed with 1,000 μL ice-cold 75% ethanol, and the samples were centrifuged again at 16,000 g for 5 min. The ethanol was removed by pipette, and a final centrifugation was conducted at 16,000 g for 1 min. The last ethanol was removed, and the samples were air dried for 20 min before the DNA was resuspended in UltraPure™ Distilled Water (Thermo Fisher Scientific). The isolations were quantified using a Qubit® dsDNA HS assay kit and a Qubit® 2.0 Fluorometer (Thermo Fisher Scientific) according the manufacturer’s instructions, and ~1 μg of DNA for each sample was shipped overnight on dry ice to Novogene Co., Ltd. (Sacramento, CA, USA) for library preparation and 150 base pair (bp) paired-end whole-genome sequencing on an Illumina NovaSeq 6000 (Illumina Inc., San Diego, CA, USA).

### Pool-seq: re-assessment of genomic mapping approaches

A previously published study assessed analytical options to determine nuclear allele frequencies from Pool-seq data for hybrid *T. californicus* by using a pool of 300 F_2_ hybrid egg sacs from a cross between the SD population and a population from Abalone Cove, California (AB: 33° 44′ 13″ N, 118° 22′ 26″ W; SD♀ x AB♂ [SDxAB]; Lima and Willett 2018). Allele frequency deviations from neutral expectations of 0.5 in F_2_ hybrids do not occur before hatch in *T. californicus* (Willett and Berkowitz 2007; Foley et al. 2013; Willett et al. 2016), establishing a clear prediction for Pool-seq data for unhatched F_2_ nauplii. Lima and Willett (2018) concluded that averaging the estimated allele frequencies from mapping a hybrid sample to the reference genomes for each population in the cross separately was the best approach to minimize the effects of mapping bias. However, there are downsides to this approach both computationally and bioinformatically relative to mapping to a single hybrid reference genome. Since these analyses were reported, an updated *T. californicus* reference genome has been published (Barreto et al. 2018), and thus in the current study we re-evaluated the potential use of a hybrid reference genome to determine nuclear allele frequencies from Pool-seq data in *T. californicus*.

Whole-genome sequencing reads were obtained for a pool of AB adults (Barreto et al. 2018) and the SDxAB F_2_ pool of unhatched nauplii from Lima & Willett (2018; NCBI SRA: SRR6391912). Adapter sequences were removed, and reads with a Phred score < 25 or a length < 50 bp were discarded. An updated population-specific genome for the AB population was created as described in Barreto et al. (2018) and Lima et al. (2019), and a hybrid reference genome for the SD and AB populations was generated from the published *T. californicus* reference genome (SD population; NCBI GenBank: GCA_007210705.1, Barreto et al. 2018) and the updated AB genome as described in Healy and Burton (2022). Note the genomes were masked such that any ‘N’ position in one genome was also ‘N’ in the other genome prior to the assembly of the hybrid reference or the read mapping described below. Single-nucleotide polymorphisms (SNPs) that displayed fixed differences between the SD and AB populations were identified following previously published protocols (Lima and Willett 2018; Lima et al. 2019; Healy and Burton 2020).

The 112,070,611 filtered reads for SDxAB nauplii were mapped to the SD reference genome, the AB reference genome and the hybrid reference genome using *BWA MEM* v0.7.12 (Li 2013) with mapping percentages of 91.87%, 92.49% and 93.32%, respectively, and alignments with a MAPQ score < 20 were discarded. For the mappings to the SD and AB genomes independently (i.e., the ‘separate mapping’), allele frequencies at the fixed SNPs between the populations were estimated for the mapping to each genome with *SAMtools* v1.14 (Danecek et al. 2021) and *PoPoolation2* v1.201 (Kofler et al. 2011) and then averaged as in Healy and Burton (2020). For the mapping to the hybrid reference genome (i.e., the ‘hybrid mapping’), the population-specific identifiers for the homologous chromosomes were removed from the alignments, and then a ‘pileup’ file was created against the masked version of the SD genome with *SAMtools* v1.14. This was possible, because population-specific genomes from resequencing in *T. californicus* have the same structure as the SD reference genome (see Barreto et al. 2018). Nuclear allele frequencies for the fixed SNPs between SD and AB with a minimum coverage of 50X and a maximum coverage of 400X were extracted from the pileup file with *PoPoolation2* v1.201.

Allele frequencies determined for individual SNPs from the separate and hybrid mappings were averaged across non-overlapping 1.5 Mb chromosomal windows of the *T. californicus* genome. Variation from the expected allele frequency of 0.5 was assessed for the windowed estimates from each mapping with Fisher’s exact tests, and variation in allele frequencies between the mappings was examined with Fisher’s exact tests and Kolmogorov-Smirnov (KS) tests as in Lima and Willett (2018), Lima et al. (2019) and Healy and Burton (2020). Finally, associations between the allele frequencies from the two mappings were tested with Pearson correlations both for the individual SNPs and for the chromosomal windows.

### Pool-seq: data analysis and statistics for fast- and slow-developing F_2_ hybrids

Based on the SDxAB F_2_ nauplii re-assessment (see Results), our sequencing reads for the SDxBR, BRxSD, SCxPE and PExSC hybrids were mapped to their corresponding hybrid reference genomes (i.e., SD+BR or SC+PE). Prior to assembly of the hybrid references as described in Healy and Burton (2022), population-specific genomes for BR and PE were created using the methods of Barreto et al. (2018) and Lima et al. (2019) from whole-genome sequencing reads for a pool of BR adults (Barreto et al. 2018) and a pool of 200 PE adults from the laboratory cultures in the current study. DNA for the PE pool was isolated and sequenced as describe above for the F_2_ hybrid pools, and 148,176,596 paired-end reads were obtained. Previously published sequences were available for the SD reference genome, a SC population-specific genome (Healy and Burton 2020), and for the SD (NCBI GenBank: DQ913891.2; Burton et al. 2007), BR (Barreto et al. 2018) and SC (NCBI GenBank: DQ917374.1; Burton et al. 2007) mitochondrial genomes. A mitochondrial sequence for the PE population was generated by mapping the PE pool of sequencing reads to the SC mitochondrial sequence and calling a consensus PE sequence in *CLC Genomic Workbench* v21 (Qiagen, Germantown, MD, USA). After assembly of the hybrid reference genomes, the F_2_ hybrid samples were mapped to the corresponding hybrid genome with *BWA MEM* v0.7.12 (Li 2013) and nuclear allele frequencies at SNPs fixed between the SD and BR or SC and PE populations were determined with *SAMtools* v1.14 (Danecek et al. 2021) and *PoPoolation2* v1.201 (Kofler et al. 2011) as outlined for the SDxAB hybrid reference mapping above. The SNPs that were fixed between the population pairs were identified following the methods of Lima and Willett (2018), Lima et al. (2019) and Healy and Burton (2020), and allele frequencies were determined for the common set of SNPs with a minimum coverage of 50X and a maximum coverage of 400X across all the pools for each pair of reciprocal crosses.

All statistical analyses were conducted in *R* v4.2.0 (R Core Team 2022) with α = 0.05 unless otherwise noted, and tests relying on count data utilized effective counts, which adjust the counts for the random chromosomal re-sampling that occurs in Pool-seq data (Wiberg et al. 2017). Allele frequencies were compared between pools of fast- and slow-developers within each cross, between fast developers from each reciprocal cross and between slow developers from each reciprocal cross. Differences between pools were assessed by (1) Fisher’s exact tests for individual SNPs followed by false-discovery rate corrections with the Benjamini-Hochberg method (Benjamini and Hochberg 1995), and (2) KS tests for the windowed allele frequencies across each chromosome with a Bonferroni correction of α = 4.17 × 10^−3^ (see Lima et al. 2019, Healy and Burton 2020). Additionally, differences from neutral expectations (i.e., 0.5) favouring the same allele in the F_2_ adult pools were examined by comparing 10^th^ and 90^th^ percentiles of the windowed frequencies for each chromosome to the 10^th^ and 90^th^ percentiles of windowed allele frequencies in the SDxAB F_2_ nauplii, which reflect neutral variation around a value of 0.5 (i.e., experimental estimate error), as in Lima et al. (2019) and Healy and Burton (2020).

## Results

### Allele frequency estimation using a hybrid reference genome in F_2_ nauplii

Allele frequencies were determined for 2,977,543 fixed SNPs between the SD and AB populations in the unhatched F_2_ SDxAB nauplii, and the coverages of the SNP loci were 97 ± 28X and 100 ± 29X, μ ± σ, in the separate and hybrid mappings, respectively. The average SD allele frequencies across all the SNPs were 0.498 ± 0.091, μ ± σ, in the separate mapping and 0.496 ± 0.090 in the hybrid mapping, and there was little variation from a frequency of 0.5 across the genome in either mapping (Fig. 1a,b). Moreover, there were no significant differences from 0.5 for any chromosomal window in either mapping (*p* ≤ 0.45), and no differences between the mapping approaches (Fisher’s exact test FDR-adjusted *p* ≤ 0.99; KS test *p* ≤ 0.17). The allele frequencies estimated with either mapping approach were also highly correlated either for the chromosomal windows (*p* < 2.2 × 10^−16^; r^2^ = 0.99; Fig. 1c), or for individual SNPs (*p* < 2.2 × 10^−16^; r^2^ = 0.94), and in both cases the lines of best fit between the two mapping approaches were similar to a 1:1 line (slope = 1.00 or 0.97, and intercept = −0.004 or 0.015 for the windows or individual SNPs, respectively). These results clearly demonstrate that mapping to a hybrid reference is an appropriate approach to estimate allele frequencies from Pool-seq data in *T. californicus*.

**Figure 1.**
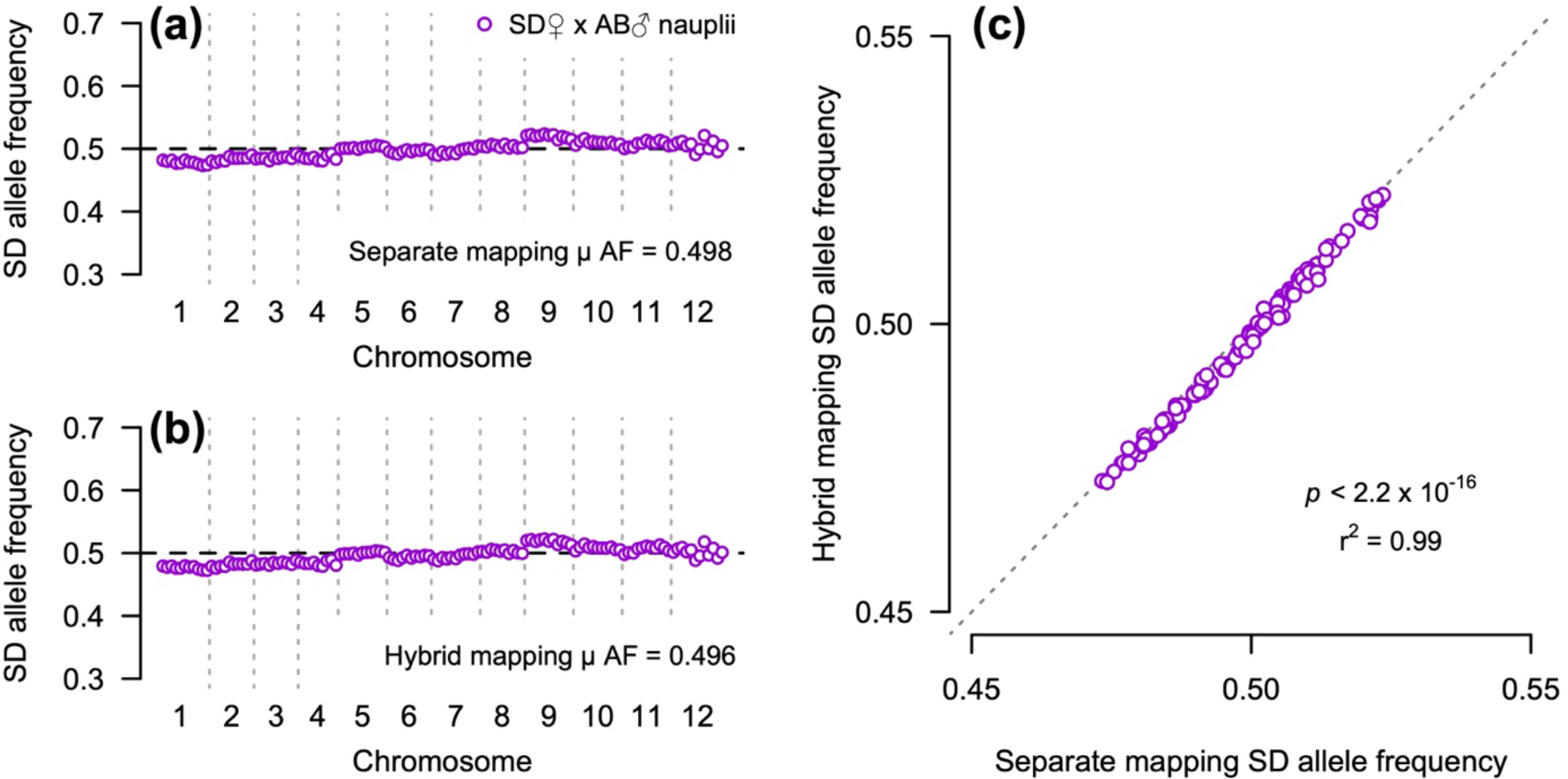
Comparisons of nuclear allele frequencies for unhatched F_2_ SDxAB hybrid nauplii from Lima & Willett (2018) after mapping to separate population reference genomes or to a hybrid reference genome. SD allele frequencies are shown for 1.5 Mb chromosomal windows across the twelve *T. californicus* nuclear chromosomes (a: separate mapping, μ = 0.498; b: hybrid mapping, μ = 0.496; no statistical differences from the expected frequency of 0.5 or between the two mappings). Panel (c) displays a significant correlation between windowed allele frequencies from the hybrid mapping against the equivalent frequencies for the separate mapping (*p*-value and r^2^ for windows displayed on the graph; *p* < 2.2 × 10^−16^ and r^2^ = 0.94 for correlation using frequencies for individual SNPs; dotted grey line – 1:1 line).

### Allele frequency variation between fast- and slow-developing F_2_ hybrids

Allele frequencies for 261,237 fixed nuclear SNPs between the SD and BR populations were scored for our SDxBR and BRxSD F_2_ hybrids. In hybrids with the SD mitochondrial genotype (i.e., SDxBR), significant differences between fast and slow developers at individual loci were detected on every chromosome (Fig. 2a,b), reflecting the highly variable nature of frequency estimates for individual SNPs from Pool-seq (Kofler et al. 2011; Lima et al. 2019), but there were clear concentrations of significant SNPs on chromosomes 1, 2, 5, 6 and 7 (43-165 per chromosome). Consistent with these patterns in general, differences between the fast- and slow-developing copepodids were detected at the chromosomal level for chromosomes 2, 5, 6 and 7 by KS tests (*p* ≤ 6.5 × 10^−4^). Additionally, allele frequency deviations from 0.5 that suggested effects of only nuclear loci on hybrid allele frequencies were observed for chromosomes 3 and 7. In hybrids with the BR mitochondrial genotype (i.e., BRxSD), again, significant differences between fast and slow developers were identified for individual SNPs on every chromosome (Fig. 2c,d), but for this cross there were no obvious concentrations of significant SNPs on any chromosome (maximum of 24 on chromosome 7). This reduced variation between fast- and slow-developing copepodids compared to the variation in the SDxBR cross was also observed at the chromosomal level, as only chromosome 6 displayed a significant difference between the BRxSD fast and slow developers (KS test *p* = 1.1 × 10^−5^). In addition, the only chromosome with allele frequency patterns consistent with potential effects of only nuclear loci was chromosome 7.

**Figure 2.**
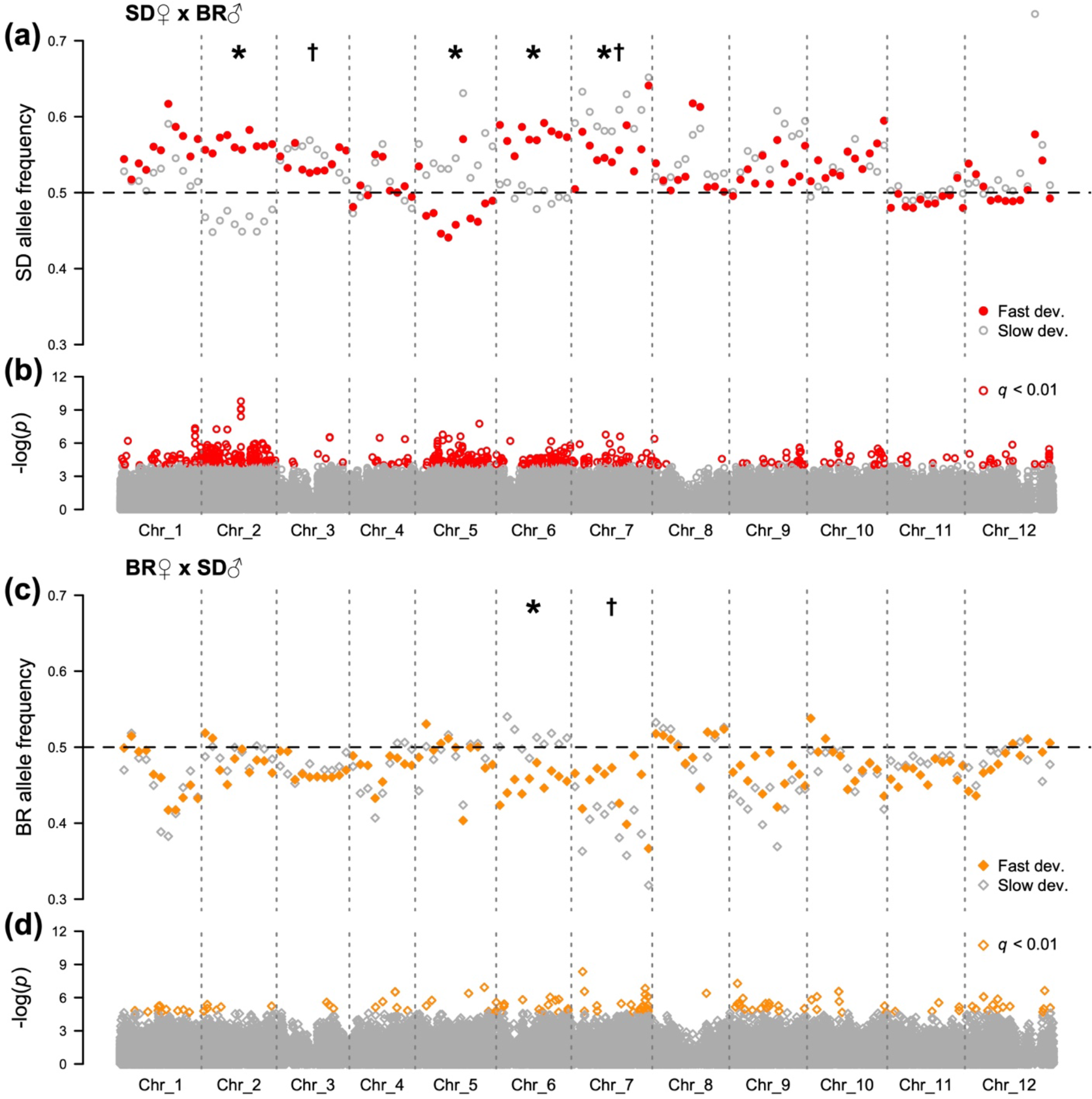
Nuclear allele frequencies for 1.5 Mb chromosomal windows in fast- and slow-developing F_2_ hybrids between the SD and BR populations (a: SDxBR, SD allele frequencies, fast developers – filled red circles, slow developers – empty grey circles; c: BRxSD, BR allele frequencies, fast developers – filled orange diamonds, slow developers – empty grey diamonds). Panels (b) and (d) display statistical results for individual loci (significant loci: b – empty red circles, d – empty orange diamonds). Asterisks indicate chromosomes with significant differences between reciprocal crosses based on KS tests, and daggers indicate chromosomes with patterns consistent with nuclear-only effects.

There was generally less variation in nuclear allele frequencies between fast- and slow-developing F_2_ hybrids from the crosses between the SC and PE populations than from the crosses between the SD and BR populations. In the SCxPE and PExSC crosses, allele frequencies were determined at 52,502 nuclear SNPs, and relatively few individual SNPs had frequencies that were significantly different between fast and slow developers (totals of 3 or 24 across all chromosomes in SCxPE or PExSC, respectively; Fig. 3). However, significant differences at the chromosomal level were detected for chromosome 9 in hybrids with the SC mitochondrial genotype (i.e., SCxPE; KS test *p* = 6.5 × 10^−4^) and for chromosomes 3, 5 and 7 in hybrids with the PE mitochondrial genotypes (i.e., PExSC; KS test *p* ≤ 2.2 × 10^−4^). In contrast, no patterns of allele frequency variation suggesting the effects of only nuclear loci were evident in either reciprocal cross between the SC and PE populations.

**Figure 3.**
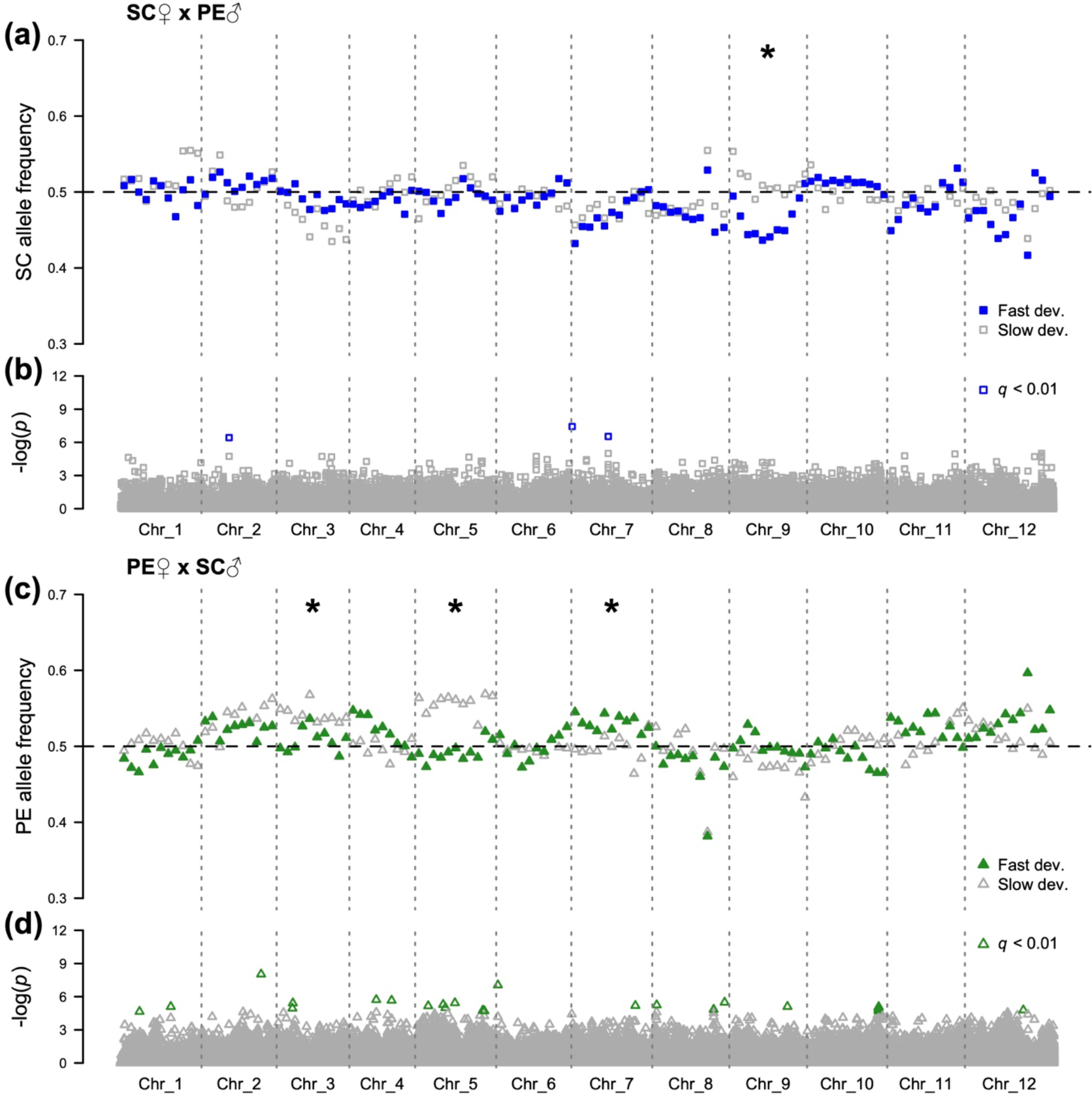
Nuclear allele frequencies for 1.5 Mb chromosomal windows in fast- and slow-developing F_2_ hybrids between the SC and PE populations (a: SCxPE, SC allele frequencies, fast developers – filled blue squares, slow developers – empty grey squares; c: PExSC, PE allele frequencies, fast developers – filled green triangles, slow developers – empty grey triangles). Panels (b) and (d) display statistical results for individual loci (significant loci: b – empty blue squares, d – empty green triangles). Asterisks indicate chromosomes with significant differences between reciprocal crosses based on KS tests.

### Allele frequency variation between reciprocal F_2_ hybrids

In previous studies in *T. californicus*, the clearest signatures of intergenomic coevolution and genetic incompatibilities in F_2_ hybrids were demonstrated by differences in allele frequencies between reciprocal hybrids, particularly between fast developers (Healy and Burton 2020; Han and Barreto 2021). Consequently, beyond the comparisons between fast- and slow-developing hybrids within crosses, we also analyzed differences between SDxBR and BRxSD hybrids, and between SCxPE and PExSC hybrids in the current study.

In fast-developing hybrids, individual SNPs with significant differences in allele frequencies between the SDxBR and BRxSD crosses, or between the SCxPE and PExSC crosses were detected on all chromosomes (Fig. 4), but there were generally no clear concentrations of significant SNPs on any chromosome (maximum of 37 on chromosome 9 for crosses between SD and BR or of 22 on chromosome 8 for crosses between SC and PE). At the chromosomal level, differences between SDxBR and BRxSD were found for three chromosomes (chromosomes 2, 6 and 11; KS test *p* ≤ 2.1 × 10^−3^), whereas only two chromosomes displayed differences between SCxPE and PExSC (chromosomes 2 and 8; KS test *p* ≤ 2.2 × 10^−4^). Variation in allele frequencies consistent with nuclear-only effects were observed on chromosomes 6 and 7 in fast developers, but only in hybrids between the SD and BR populations.

**Figure 4.**
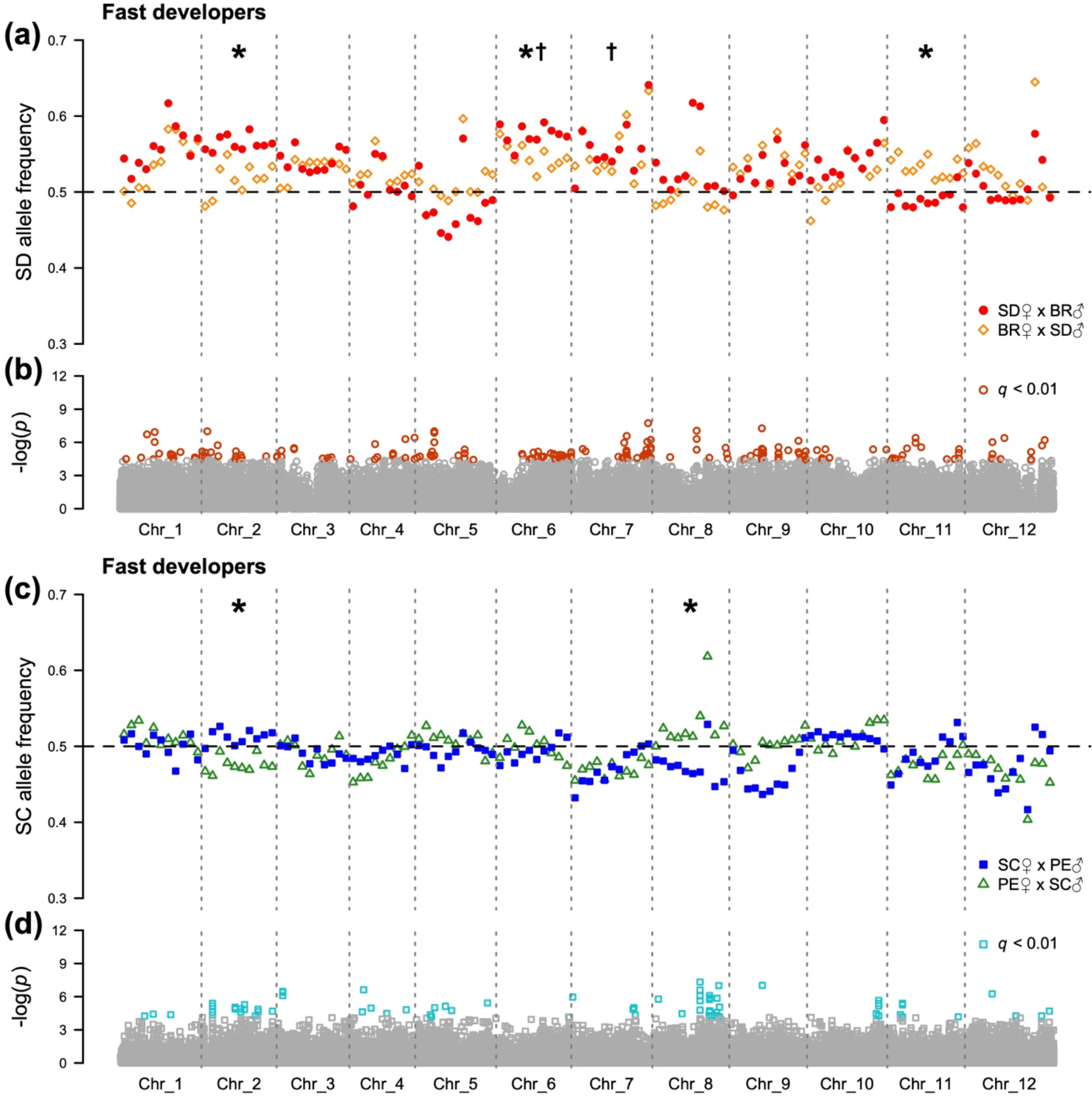
Nuclear allele frequencies for 1.5 Mb chromosomal windows in fast-developing F_2_ hybrids (a: SD allele frequencies, SDxBR – filled red circles, BRxSD – empty orange diamonds; c: SC allele frequencies, SCxPE – filled blue squares, PExSC – empty green triangles). Panels (b) and (d) display statistical results for individual loci (significant loci: b – empty dark orange circles, d – empty turquoise squares). Asterisks indicate chromosomes with significant differences between reciprocal crosses based on KS tests, and daggers indicate chromosomes with patterns consistent with nuclear-only effects.

As observed in the fast-developing copepodids, allele frequencies at individual SNPs in slow-developing hybrids were significantly different between the SDxBR and BRxSD crosses, and the SCxPE and PExSC crosses for loci on all chromosomes (Fig. 5). Again, there were no clear concentrations of significant loci on any chromosome (maximum of 19 on chromosome 5 for crosses between SD and BR or of 22 on chromosome 8 for crosses between SC and PE). KS tests detected differences at the chromosome level for chromosomes 2 and 11 (*p* ≤ 1.1 × 10^−5^) between SDxBR and BRxSD, and for chromosomes 2 and 5 between SCxPE and PExSC (*p* ≤ 2.1 × 10^−3^). No allele frequency patterns consistent with effects of only nuclear loci were observed for slow developers from crosses between the SC and PE populations, whereas relatively large signatures of nuclear-only effects were displayed on chromosomes 7 and 9 in slow developers from the SDxBR and BRxSD crosses.

**Figure 5.**
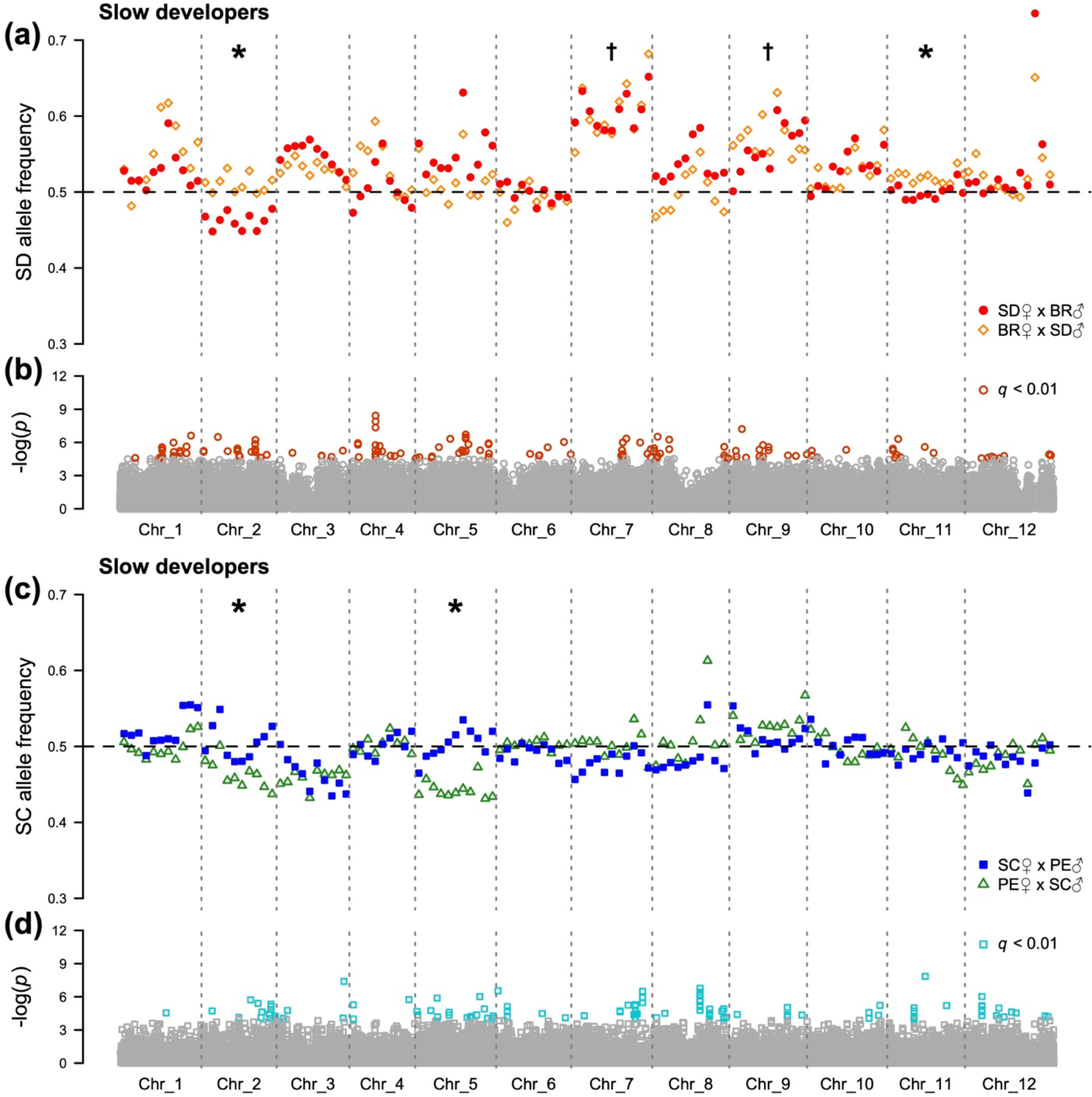
Nuclear allele frequencies for 1.5 Mb chromosomal windows in slow-developing F_2_ hybrids (a: SD allele frequencies, SDxBR – filled red circles, BRxSD – empty orange diamonds; c: SC allele frequencies, SCxPE – filled blue squares, PExSC – empty green triangles). Panels (b) and (d) display statistical results for individual loci (significant loci: b – empty dark orange circles, c – empty turquoise squares). Asterisks indicate chromosomes with significant differences between reciprocal crosses based on KS tests, and daggers indicate chromosomes with patterns consistent with nuclear-only effects.

## Discussion

The role of mitonuclear incompatibilities in eukaryotic hybrid breakdown remains unclear as loss of fitness has been associated with incompatible mitonuclear genotypes in some species (e.g., Meiklejohn et al. 2013), whereas introgression of mitochondrial DNA is often observed in natural zones of admixture between populations or species (e.g., Toews and Brelsford 2012). Strong selection associated with mitonuclear interactions has previously been demonstrated using fast-developing inter-population hybrids of *T. californicus*; however, these hybrids were from crosses between highly divergent (> 20% mtDNA sequence divergence) and geographically distant (> 600 km) populations (Healy and Burton 2020; Han and Barreto 2021). To assess if these high levels of divergence are necessary to observe strong effects of mitonuclear incompatibilities, the current study uses two pairs of less divergent and less geographically distant populations; SC and PE are ~50 km apart and the nucleotide sequences of their mitochondrial genomes are only 1.5% divergent (current study), and SD and BR are separated by ~8 km and divergence between their mitochondrial genotypes is 9.6% (Barreto et al. 2018). Despite these reductions in inter-population divergence and distance, allele frequency deviations consistent with effects of mitonuclear interactions and with maintenance of coevolved genotypes were observed in both crosses, particularly in fast-developing (high-fitness) F_2_ hybrids. Yet, although signatures of mitonuclear coevolution dominated the genomes of high-fitness hybrids from crosses between highly divergent populations, this pattern was reduced in hybrids from the low-divergence crosses. Instead, allele frequencies in F_2_ hybrids indicated not only mitonuclear effects favouring coevolved alleles, but also mitonuclear effects favouring mismatched (i.e., not coevolved) alleles and effects involving only nuclear loci. Therefore, our findings suggest that strong effects of mitonuclear incompatibilities are possible in hybrids between taxa with relatively low genetic divergence, but that these effects are components of complex suites of genetic interactions underlying hybrid breakdown overall.

Potential signatures of inter-genomic coevolution in our data are evident through excesses of nuclear alleles originating from the same population as the mitochondrial genotype in reciprocal fast-developing hybrids. Among the *T. californicus* crosses examined to date, the number of nuclear chromosomes displaying effects of mitonuclear coevolution tends to decrease with decreasing genetic divergence between the populations that contribute to the cross. For instance, five and seven nuclear chromosomes displayed signatures of coevolution in reciprocal fast-developing hybrids from the high-divergence crosses between the SD and SC populations, and the SD and Strawberry Hill Wayside (SH; 44° 15′ 0″ N, 124° 6′ 36″ W) populations, respectively (Healy and Burton 2020; Han and Barreto 2021). In contrast, only two chromosomes showed clear allele frequency deviations favouring coevolved nuclear alleles in crosses between SD and BR (chromosomes 2 and 6 between reciprocal fast-developing hybrids), and there were only relatively weak signatures of coevolution for one chromosome in crosses between SC and PE (chromosome 2 between both reciprocal fast- and slow-developing hybrids). However, this trend was not observed for the biases favouring mismatched genotypes. Healy and Burton (2020) found two chromosomes with biases for mismatched alleles in hybrids between SD and SC (chromosomes 6 and 9), and Han and Barreto (2021) identified one chromosome displaying this pattern in hybrids between SD and SH (chromosome 7). Similarly, in hybrids between the SD and BR populations and hybrids between the SC and PE populations, one chromosome per cross had allele frequencies consistent with mitonuclear interactions favouring mismatched genotypes (chromosome 11 between the fast- and slow-developing hybrids between SD and BR, and chromosome 8 between the fast-developing hybrids between SC and PE). This suggests that the extent of mitonuclear interactions favouring mismatched alleles is relatively independent of the divergence of the mitochondrial genome between the two *T. californicus* populations that are crossed, which is unlike the positive relationship between the effects of coevolution and inter-population divergence.

There is also not a straightforward relationship between inter-population genetic divergence and the numbers of potential effects involving only nuclear loci (i.e., where the same allele is favored in both reciprocal crosses). In the current study, no allele frequencies in the SCxPE and PExSC hybrids were consistent with nuclear-only effects, and three to four chromosomes displayed potential nuclear-only effects in the SDxBR and BRxSD crosses (chromosomes 6, 7 and 9 between the reciprocal crosses, and chromosome 3 between fast- and slow-developing SDxBR hybrids). On the other hand, hybrids from crosses between populations with high levels of genetic divergence demonstrated nuclear-only effects (potentially nuclear-nuclear interactions; Lima et al. 2019) on either one or two chromosomes (chromosome 3 in hybrids between SD and SH; Han and Barreto 2021, and chromosomes 8 and 11 in hybrids between SD and SC; Healy and Burton 2020). Therefore, the ratios of mitonuclear effects consistent with coevolution to effects favouring mitonuclear mismatch to effects of nuclear variation alone were higher in the hybrids from high divergence crosses (5:2:2 or 7:1:1 in hybrids between SD and SC or SH, respectively) than in the hybrids from the low divergence crosses (2:1:4 in hybrids between SD and BR, and 1:1:0 in hybrids between SC and PE). These decreases in the relative proportion of positive effects of coevolution in the low-divergence crosses are similar to the results of studies in *T. californicus* using crosses between populations with higher levels of genetic divergence (e.g., between SD and SC) in which the F_2_ hybrids were not sorted into high- or low-fitness groups (Edmands et al. 2009; Pritchard et al. 2011; Foley et al. 2013; Willett et al. 2016; Lima et al. 2019). Taken together, these results indicate that mitonuclear interactions affect developmental rate (i.e., fitness) and nuclear allele frequencies in F_2_ hybrids even in crosses between populations with relatively low genetic divergence. However, the relative influence of coevolved mitonuclear interactions compared to other genetic effects or interactions is reduced in crosses between low-divergence populations.

The majority of mitonuclear interactions were observed consistently across entire chromosomes in the current study, which is expected given that large regions of chromosomes are inherited together in F_2_ hybrids due to only one opportunity for inter-population recombination (Lima and Willett 2018; Lima et al. 2019), which occurs only in male *T. californicus* (Burton et al. 1981). Although there were not clear biases for chromosomes displaying effects of coevolution in the crosses between populations of relatively low genetic divergence in the current study, substantial allele frequency deviations favouring coevolved nuclear alleles were still evident in these crosses. For example, in the SDxBR hybrids carrying SD mitochondrial DNA, fast developers had excess SD alleles on chromosome 2, whereas slow developers had excess BR alleles. Furthermore, comparisons among fast developers in SDxBR and BRxSD hybrids, and in SCxPE and PExSC hybrids also demonstrated differences consistent with coevolution on chromosome 2. Allele frequency deviations favouring coevolved alleles on this chromosome reached up to 0.083 in crosses between the SD and BR populations and up to 0.039 in crosses between the SC and PE populations. These differences may appear modest, but in *T. californicus* there is little evidence of selection against heterozygotes in F_2_ hybrids (Pritchard et al. 2011; Foley et al. 2013), and consequently the chromosome 2 allele frequency deviations reported here suggest that hybrids with homozygous mitonuclear mismatches on this chromosome are underrepresented by up to 57% or 29% in the crosses between the SD and BR populations or the SC and PE populations, respectively (compare to underrepresentation of homozygous mismatches by 87% on chromosome 2 in hybrids between SD and SC; Healy and Burton 2020). Chromosome 2 is the only nuclear chromosome to be consistently associated with allele frequency biases favouring coevolved alleles in fast-developing F_2_ *T. californicus* hybrids in all four of the inter-population crosses examined to date (Healy and Burton 2020; Han and Barreto 2021; current study). Interestingly, the magnitude of the deviations favouring coevolved alleles generally follows the divergence of the mitochondrial genome sequence between the two populations contributing to the cross (e.g., 0.138 and 21.7% for SD to SC, 0.096 and 21.1% for SD to SH, 0.083 and 9.6% for SD to BR, and 0.039 and 1.5% for SC to PE), although there are too few observations available to assess their statistical significance. Alternatively, this may indicate that multiple loci on this chromosome develop coevolved interactions as divergence progresses (see Willett et al. 2016 for evidence for effects of multiple loci on chromosome 3), or that divergence strengthens the effects of incompatibilities over time despite the fact there is no direct selection on incompatibilities between these allopatric populations (Burton and Barreto 2012). Regardless, these results demonstrate that high levels of inter-population genetic divergence are not required for there to be strong selection favouring mitonuclear compatibility among hybrids.

Physiological research in hybrid *T. californicus* has demonstrated negative effects of mitonuclear incompatibilities on functions of the mitochondrial electron transport system (ETS; Ellison and Burton 2006, 2008b; Barreto and Burton 2013; Healy and Burton 2020; Han and Barreto 2021) and on rates of mitochondrial transcription (Ellison and Burton 2008a). In addition, elevated rates of mitochondrial ribosomal protein evolution reflect their interaction with fast-evolving mitochondrial-encoded rRNAs (Barreto and Burton 2012). These findings represent three of the four major mitochondrial functions expected to involve genes with direct inter-genomic interactions (Burton and Barreto 2012; Hill 2015, 2017; Hill et al. 2019). The remaining category of likely mitonuclear interactions is between mitochondrial-encoded tRNAs and their corresponding nuclear-encoded aminoacyl-tRNA synthetases, and there is a well-characterized incompatibility between the tRNA and aminoacyl-tRNA synthetase for tyrosine in hybrids between *D. melanogaster* and *D. simulans* that results in loss of hybrid fitness (Meiklejohn et al. 2013). No mitochondrial DNA or RNA polymerases are encoded on chromosome 2 in the nuclear genome of *T. californicus*, but proteins involved in the other three expected categories of mitonuclear interactions are produced from chromosomes 2 genes: three ETS subunits (complex I: *ndufb7* and *ndufv2*; complex III: *uqcr10*), twelve mitochondrial ribosomal proteins and three aminoacyl-tRNA synthetases. These genes are not necessarily those underlying the consistent effects of mitonuclear coevolution observed across inter-population crosses of *T. californicus*; however, they are, perhaps, the best candidate genes to underlie mitonuclear effects on chromosome 2 both from theoretical expectations and from observations of elevated rates of sequence evolution of ETS subunits, mitochondrial ribosomal proteins and aminoacyl-tRNA synthetases among populations of *T. californicus* generally (Barreto et al. 2018).

In summary, the results of the current study clearly demonstrate the complex genetic basis of variation in fitness-related traits in F_2_ hybrids, and thus of hybrid breakdown more generally. Although in comparison to previously published results in *T. californicus* substantial biases for effects of mitonuclear coevolution were reduced in F_2_ hybrids from crosses between populations with relatively low genetic divergence, effects of interactions between the genes encoded in the mitochondrial and nuclear genomes were clearly evident and were still major genetic factors contributing to variation in fitness among individual hybrids. Moreover, strong effects favouring coevolved mitonuclear genotypes were observed for at least one chromosome in the low-divergence crosses, suggesting selection acts to maintain mitonuclear compatibility, and in turn limit gene flow, even before high levels of mitochondrial divergence are achieved.

## References

Ballard, J.W.O. (2000) Comparative genomics of mitochondrial DNA in *Drosophila simulans*. J. Mol. Evol., 51, 64–75. https://doi.org/10.1007/s002390010067

Barreto, F.S. & Burton, R.S. (2012) Evidence for compensatory evolution of ribosomal proteins in response to rapid divergence of mitochondrial rRNA. Mol. Biol. Evol., 30, 310–314. https://doi.org/10.1093/molbev/mss228

Barreto, F.S. & Burton, R.S. (2013) Elevated oxidative damage is correlated with reduced fitness in interpopulation hybrids of a marine copepod. Proc. R. Soc. B, 280, 20131521. https://doi.org/10.1098/rspb.2013.1521

Barreto, F.S., Watson, E.T., Lima, T.G., Willett, C.S., Edmands, S., Li, W. & Burton, R.S. (2018) Genomic signatures of mitonuclear coevolution across populations of *Tigriopus californicus*. Nat. Ecol. Evol., 2, 1250–1257. https://doi.org/10.1038/s41559-018-0588-1

Benjamini, Y. & Hochberg, Y. (1995) Controlling the false discovery rate: a practical and powerful approach to multiple testing. J. R. Stat. Soc. Ser. B, 57, 289–300. https://doi.org/10.1111/j.2517-6161.1995.tb02031.x

Burton, R.S. (1985) Mating system of the intertidal copepod *Tigriopus californicus*. Mar. Biol., 86, 247–252. https://doi.org/10.1007/BF00397511

Burton, R.S. (1990) Hybrid breakdown in developmental time in the copepod *Tigriopus californicus*. Evolution, 44, 1814–1822. https://doi.org/10.1111/j.1558-5646.1990.tb05252.x

Burton, R.S. (2022) The role of mitonuclear incompatibilities in allopatric speciation. Cell. Mol. Life Sci., 79, 1–18. https://doi.org/10.1007/s00018-021-04059-3

Burton, R.S. & Barreto, F.S. (2012) A disproportionate role for mtDNA in Dobzhansky–Muller incompatibilities?. Mol. Ecol., 21, 4942–4957. https://doi.org/10.1111/mec.12006

Burton, R.S., Byrne, R.J. & Rawson, P.D. (2007) Three divergent mitochondrial genomes from California populations of the copepod *Tigriopus californicus*. Gene, 403, 53–59. https://doi.org/10.1016/j.gene.2007.07.026

Burton, R.S., Ellison, C.K. & Harrison, J.S. (2006) The sorry state of F_2_ hybrids: Consequences of rapid mitochondrial DNA evolution in allopatric populations. Am. Nat., 168, S14–S24. https://doi.org/10.1086/509046

Burton, R.S., Feldman, M.W. & Swisher, S.G. (1981) Linkage relationships among five enzyme-coding gene loci in the copepod *Tigriopus californicus*: A genetic confirmation of achiasmatic meiosis. Biochem. Genet., 19, 1237–1245. https://doi.org/10.1007/BF00484576

Burton, R.S. & Lee, B.N. (1994) Nuclear and mitochondrial gene genealogies and allozyme polymorphism across a major phylogeographic break in the copepod *Tigriopus californicus*. Proc. Natl. Acad. Sci. USA, 91, 5197–5201. https://doi.org/10.1073/pnas.91.11.5197

Burton, R.S., Pereira, R.J. & Barreto, F.S. (2013) Cytonuclear genomic interactions and hybrid breakdown. Annu. Rev. Ecol. Evol. Syst., 44, 281–302. https://doi.org/10.1146/annurev-ecolsys-110512-135758

Carnegie, L., Reuter, M., Fowler, K., Lane, N. & Camus, M.F. (2021) Mother's curse is pervasive across a large mitonuclear *Drosophila* panel. Evol. Lett., 5, 230–239. https://doi.org/10.1002/evl3.221

Chan, K.M. & Levin, S.A. (2005) Leaky prezygotic isolation and porous genomes: rapid introgression of maternally inherited DNA. Evolution, 59, 720–729. https://doi.org/10.1111/j.0014-3820.2005.tb01748.x

Danecek, P., Bonfield, J.K., Liddle, J., Marshall, J., Ohan, V., Pollard, M.O., Whitwham, A., Keane, T., McCarthy, S.A., Davies, R.M. & Li, H. (2021). Twelve years of SAMtools and BCFtools. GigaScience, 10, giab008. https://doi.org/10.1093/gigascience/giab008

Dowling, D.K., Friberg, U., Hailer, F. & Arnqvist, G. (2007) Intergenomic epistasis for fitness: Within-population interactions between cytoplasmic and nuclear genes in *Drosophila melanogaster*. Genetics, 175, 235–244. https://doi.org/10.1534/genetics.105.052050

Edmands, S. (2001) Phylogeography of the intertidal copepod *Tigriopus californicus* reveals substantially reduced population differentiation at northern latitudes. Mol. Ecol., 10, 1743–1750. https://doi.org/10.1046/j.0962-1083.2001.01306.x

Edmands, S. & Burton, R.S. (1999) Cytochrome *c* oxidase activity in interpopulation hybrids of a marine copepod: A test for nuclear‐nuclear or nuclear‐cytoplasmic coadaptation. Evolution, 53, 1972–1978. https://doi.org/10.1111/j.1558-5646.1999.tb04578.x

Edmands, S., Northrup, S.L. & Hwang, A.S. (2009) Maladapted gene complexes within populations of the intertidal copepod *Tigriopus californicus*?. Evolution, 63, 2184–2192. https://doi.org/10.1111/j.1558-5646.2009.00689.x

Ellison, C.K. & Burton, R.S. (2006) Disruption of mitochondrial function in interpopulation hybrids of *Tigriopus californicus*. Evolution, 60, 1382–1391. https://doi.org/10.1111/j.0014-3820.2006.tb01217.x

Ellison, C.K. & Burton, R.S. (2008a) Genotype-dependent variation of mitochondrial transcriptional profiles in interpopulation hybrids. Proc. Natl. Acad. Sci. USA, 105, 15831–15836. https://doi.org/10.1073/pnas.0804253105

Ellison, C.K. & Burton, R.S. (2008b) Interpopulation hybrid breakdown maps to the mitochondrial genome. Evolution, 62, 631–638. https://doi.org/10.1111/j.1558-5646.2007.00305.x

Eyre-Walker, A. (2017) Mitochondrial replacement therapy: Are mito-nuclear interactions likely to be a problem?. Genetics, 205, 1365–1372. https://doi.org/10.1534/genetics.116.196436

Foley, B.R., Rose, C.G., Rundle, D.E., Leong, W. & Edmands, S. (2013) Postzygotic isolation involves strong mitochondrial and sex-specific effects in *Tigriopus californicus*, a species lacking heteromorphic sex chromosomes. Heredity, 111, 391–401. https://doi.org/10.1038/hdy.2013.61

Gershoni, M., Templeton, A.R. & Mishmar, D. (2009) Mitochondrial bioenergetics as a major motive force of speciation. Bioessays, 31, 642–650. https://doi.org/10.1002/bies.200800139

Han, K.L. & Barreto, F.S. (2021) Pervasive mitonuclear coadaptation underlies fast development in interpopulation hybrids of a marine crustacean. Genome Biol. Evol., 13, evab004. https://doi.org/10.1093/gbe/evab004

Harrison, J.S. & Burton, R.S. (2006) Tracing hybrid incompatibilities to single amino acid substitutions. Mol. Biol. Evol., 23, 559–564. https://doi.org/10.1093/molbev/msj058

Healy, T.M. & Burton, R.S. (2020) Strong selective effects of mitochondrial DNA on the nuclear genome. Proc. Natl. Acad. Sci. USA, 117, 6616–6621. https://doi.org/10.1073/pnas.1910141117

Healy, T.M. & Burton, R.S. (2022) Differential gene expression and mitonuclear incompatibilities in fast- and slow-developing inter-population *Tigriopus californicus* hybrids. bioRxiv. https://doi.org/10.1101/2022.09.09.507197

Hill, G.E. (2015) Mitonuclear ecology. Mol. Biol. Evol., 32, 1917–1927. https://doi.org/10.1093/molbev/msv104

Hill, G.E. (2016) Mitonuclear coevolution as the genesis of speciation and the mitochondrial DNA barcode gap. Ecol. Evol., 6, 5831–5842. https://doi.org/10.1002/ece3.2338

Hill, G.E. (2017) The mitonuclear compatibility species concept. Auk, 134, 393–409. https://doi.org/10.1642/AUK-16-201.1

Hill, G.E. (2019) Reconciling the mitonuclear compatibility species concept with rampant mitochondrial introgression. Integr. Comp. Biol., 59, 912–924. https://doi.org/10.1093/icb/icz019

Hill, G.E., Havird, J.C., Sloan, D.B., Burton, R.S., Greening, C. & Dowling, D.K. (2019) Assessing the fitness consequences of mitonuclear interactions in natural populations. Biol. Rev., 94, 1089–1104. https://doi.org/10.1111/brv.12493

Hoekstra, L.A., Siddiq, M.A. & Montooth, K.L. (2013) Pleiotropic effects of a mitochondrial – nuclear incompatibility depend upon the accelerating effect of temperature in *Drosophila*. Genetics, 195, 1129–1139. https://doi.org/10.1534/genetics.113.154914

Kofler, R., Pandey, R.V. & Schlötterer, C. (2011) PoPoolation2: Identifying differentiation between populations using sequencing of pooled DNA samples (Pool-Seq). Bioinform., 27, 3435–3436. https://doi.org/10.1093/bioinformatics/btr589

Lane, N. (2005) Power, sex, suicide: mitochondria and the meaning of life. Oxford University Press, Oxford, United Kingdom.

Li, H. (2013) Aligning sequence reads, clone sequences and assembly contigs with BWA-MEM. arXiv, arXiv:1303.3997. https://doi.org/10.48550/arXiv.1303.3997

Lima, T.G., Burton, R.S. & Willett, C.S. (2019) Genomic scans reveal multiple mito‐nuclear incompatibilities in population crosses of the copepod *Tigriopus californicus*. Evolution, 73, 609–620. https://doi.org/10.1111/evo.13690

Lima, T.G. & Willett, C.S. (2018) Using Pool-seq to search for genomic regions affected by hybrid inviability in the copepod *T. californicus*. J. Hered., 109, 469–476. https://doi.org/10.1093/jhered/esx115

Lynch, M. (1997) Mutation accumulation in nuclear, organelle, and prokaryotic transfer RNA genes. Mol. Biol. Evol., 14, 914–925. https://doi.org/10.1093/oxfordjournals.molbev.a025834

Meiklejohn, C.D., Holmbeck, M.A., Siddiq, M.A., Abt, D.N., Rand, D.M. & Montooth, K.L. (2013) An incompatibility between a mitochondrial tRNA and its nuclear-encoded tRNA synthetase compromises development and fitness in *Drosophila*. PLoS Genet., 9, e1003238. https://doi.org/10.1371/journal.pgen.1003238

Pereira, R.J., Barreto, F.S., Pierce, N.T., Carneiro, M. & Burton, R.S. (2016) Transcriptome‐wide patterns of divergence during allopatric evolution. Mol. Ecol., 25, 1478–1493. https://doi.org/10.1111/mec.13579

Pereira, R.J., Lima, T.G., Pierce‐Ward, N.T., Chao, L. & Burton, R.S. (2021) Recovery from hybrid breakdown reveals a complex genetic architecture of mitonuclear incompatibilities. Mol. Ecol., 30, 6403–6416. https://doi.org/10.1111/mec.15985

Peterson, D.L., Kubow, K.B., Connolly, M.J., Kaplan, L.R., Wetkowski, M.M., Leong, W., Phillips, B.C. & Edmands, S. (2013) Reproductive and phylogenetic divergence of tidepool copepod populations across a narrow geographical boundary in Baja California. J. of Biogeogr., 40, 1664–1675. https://doi.org/10.1111/jbi.12107

Pritchard, V.L., Dimond, L., Harrison, J.S., Velázquez, C.C.S., Zieba, J.T., Burton, R.S. & Edmands, S. (2011) Interpopulation hybridization results in widespread viability selection across the genome in *Tigriopus californicus*. BMC Genet., 12, 1–13. https://doi.org/10.1186/1471-2156-12-54

R Core Team. (2022) R: A language and environment for statistical computing. R Foundation for Statistical Computing, Vienna, Austria. https://www.R-project.org/.

Raisuddin, S., Kwok, K.W., Leung, K.M., Schlenk, D. & Lee, J.S. (2007) The copepod *Tigriopus*: A promising marine model organism for ecotoxicology and environmental genomics. Aquat. Toxicol., 83, 161–173. https://doi.org/10.1016/j.aquatox.2007.04.005

Rand, D.M., Haney, R.A. & Fry, A.J. (2004) Cytonuclear coevolution: The genomics of cooperation. Trends Ecol. Evol., 19, 645–653. https://doi.org/10.1016/j.tree.2004.10.003

Rand, D.M., Mossman, J.A., Spierer, A.N. & Santiago, J.A. (2022) Mitochondria as environments for the nuclear genome in *Drosophila*: Mitonuclear G×G×E. J. Hered., 113, 37–47. https://doi.org/10.1093/jhered/esab066

Sambrook, J. & Russell, D.W. (2006) Purification of nucleic acids by extraction with phenol: chloroform. Cold Spring Harb. Protoc., 2006, pdb–prot4455. https://doi.org/10.1101/pdb.prot4455

Sloan, D.B., Havird, J.C. & Sharbrough, J. (2017) The on‐again, off‐again relationship between mitochondrial genomes and species boundaries. Mol. Ecol., 26, 2212–2236. https://doi.org/10.1111/mec.13959

Toews, D.P. & Brelsford, A. (2012) The biogeography of mitochondrial and nuclear discordance in animals. Mol. Ecol., 21, 3907–3930. https://doi.org/10.1111/j.1365-294X.2012.05664.x

Wallace, D.C. (2010) Mitochondrial DNA mutations in disease and aging. Environ. Mol. Mutagen., 51, 440–450. https://doi.org/10.1002/em.20586

Wiberg, R.A.W., Gaggiotti, O.E., Morrissey, M.B. & Ritchie, M.G. (2017) Identifying consistent allele frequency differences in studies of stratified populations. Methods Ecol. Evol., 8, 1899–1909. https://doi.org/10.1111/2041-210X.12810

Willett, C.S. & Berkowitz, J.N. (2007) Viability effects and not meoitic drive cause dramatic departures from Mendelian inheritance for malic enzyme in hybrids of *Tigriopus californicus* populations. J. Evol. Biol., 20, 1196–1205. https://doi.org/10.1111/j.1420-9101.2006.01281.x

Willett, C.S., Lima, T.G., Kovaleva, I. & Hatfield, L. (2016) Chromosome-wide impacts on the expression of incompatibilities in hybrids of *Tigriopus californicus*. G3-Genes Genom. Genet., 6, 1739–1749. https://doi.org/10.1534/g3.116.028050

